# Invasiveness Screening Kit (ISK) v3: an integrated multilingual decision-support platform for non-native species risk identification

**DOI:** 10.64898/2026.07.24.740490

**Authors:** Lorenzo Vilizzi, Aisha Al-Marhoun

## Abstract

Risk identification of non-native species is widely used to support prioritization, early warning, horizon scanning and management decisions. Over the past two decades, the Invasiveness Screening Kit (ISK) framework of decision-support tools has developed through aquatic, terrestrial animal and terrestrial plant applications, but until now these toolkits have been distributed and used as separate software environments. ISK v3 provides a single integrated Microsoft Excel/Visual Basic for Applications platform for the Aquatic Species Invasiveness Screening Kit (AS-ISK), Terrestrial Animal Species Invasiveness Screening Kit (TAS-ISK) and Terrestrial Plant Species Invasiveness Screening Kit (TPS-ISK). It preserves the established questionnaires, scoring logic and multilingual implementation of the three toolkits within a single integrated interface, while introducing improved harmonized database workflows, screening record management, taxonomic verification, threshold handling with built-in calibration, risk summaries, reporting, merging and export functions as well as controlled access to separate Excel instances. The platform supports 31 languages, toolkit-specific databases, conversion of compatible v2 databases, import of legacy aquatic first-generation ISK databases into AS-ISK, online verification through major taxonomic and biodiversity data resources, and use of a global *a priori* categorization dataset for calibration. ISK v3 preserves the reproducible screening structure of the three toolkits while improving consistency, transparency and traceability in non-native species risk identification and reducing fragmentation across toolkits. It provides a common platform for researchers, managers and institutions applying risk identification workflows across aquatic, terrestrial animal and terrestrial plant taxa.

## Introduction

Biological invasions represent a major component of global environmental change (Blackburn et al. 2011; Early et al. 2016), with non-native species (*sensu* Soto et al. 2024; Vilizzi et al. 2025b, 2026b) able to adversely affect biodiversity, ecosystem processes and socio-economic systems (Haubrock et al. 2026a). Because management interventions become increasingly difficult, costly and uncertain once a species is established and spreading in its introduced range (Leung et al. 2002; Hulme 2009; Haubrock et al. 2026b), prevention and early prioritization are central to biosecurity planning (Roy et al. 2019, 2024). Within invasion risk analysis, risk identification, also referred to as risk screening, represents the initial step in determining whether a non-native species may be introduced, establish, spread and ultimately cause adverse ecological or socio-economic effects in a defined risk assessment area. This step differs from follow-up full risk assessment, which evaluates the likelihood and magnitude of risk in greater detail, and from final risk management, which concerns the selection and implementation of appropriate responses (see Vilizzi et al. 2026b).

Decision-support tools have therefore become important instruments for scientists, environmental managers and policy practitioners who need to screen potentially invasive species in a transparent, repeatable and time-efficient manner (Roy et al. 2018; Srėbalienė et al. 2019). Among these, tools derived from the Australian Weed Risk Assessment (WRA) framework (Pheloung et al. 1999; Gordon et al. 2008) have been particularly influential because they combine structured questionnaires, evidence-based responses and semi-quantitative scoring to support the ranking of species into low, medium and high risk. The Invasiveness Screening Kit (ISK) framework of decision-support tools developed from this tradition, initially through the first-generation ISK tools for aquatic species, beginning with the freshwater Fish Invasiveness Screening Kit (FISK: Copp et al. 2005a, 2005b; Lawson et al. 2013; Vilizzi et al. 2019), including its Spanish version (S-FISK), and followed by other taxon-specific toolkits comprising the Amphibian Invasiveness Screening Kit (Amph-ISK), the Freshwater Invertebrate Screening Kit (FI-ISK), the Marine Fish Invasiveness Screening Kit (MFISK) and the Marine Invertebrate Invasiveness Screening Kit (MI-ISK) (Copp 2013). These first- generation aquatic tools were superseded by the second-generation ISK tools including the Aquatic Species Invasiveness Screening Kit (AS-ISK: Copp et al. 2016, 2021), which expanded the range of aquatic organisms to include all taxonomic groups from freshwater, brackish and marine habitats, and the more recently developed Terrestrial Animal Species Invasiveness Screening Kit (TAS-ISK: Vilizzi et al. 2022c) and Terrestrial Plant Species Invasiveness Screening Kit (TPS-ISK: Vilizzi et al. 2025a) (Fig. 1).

**Figure 1.**
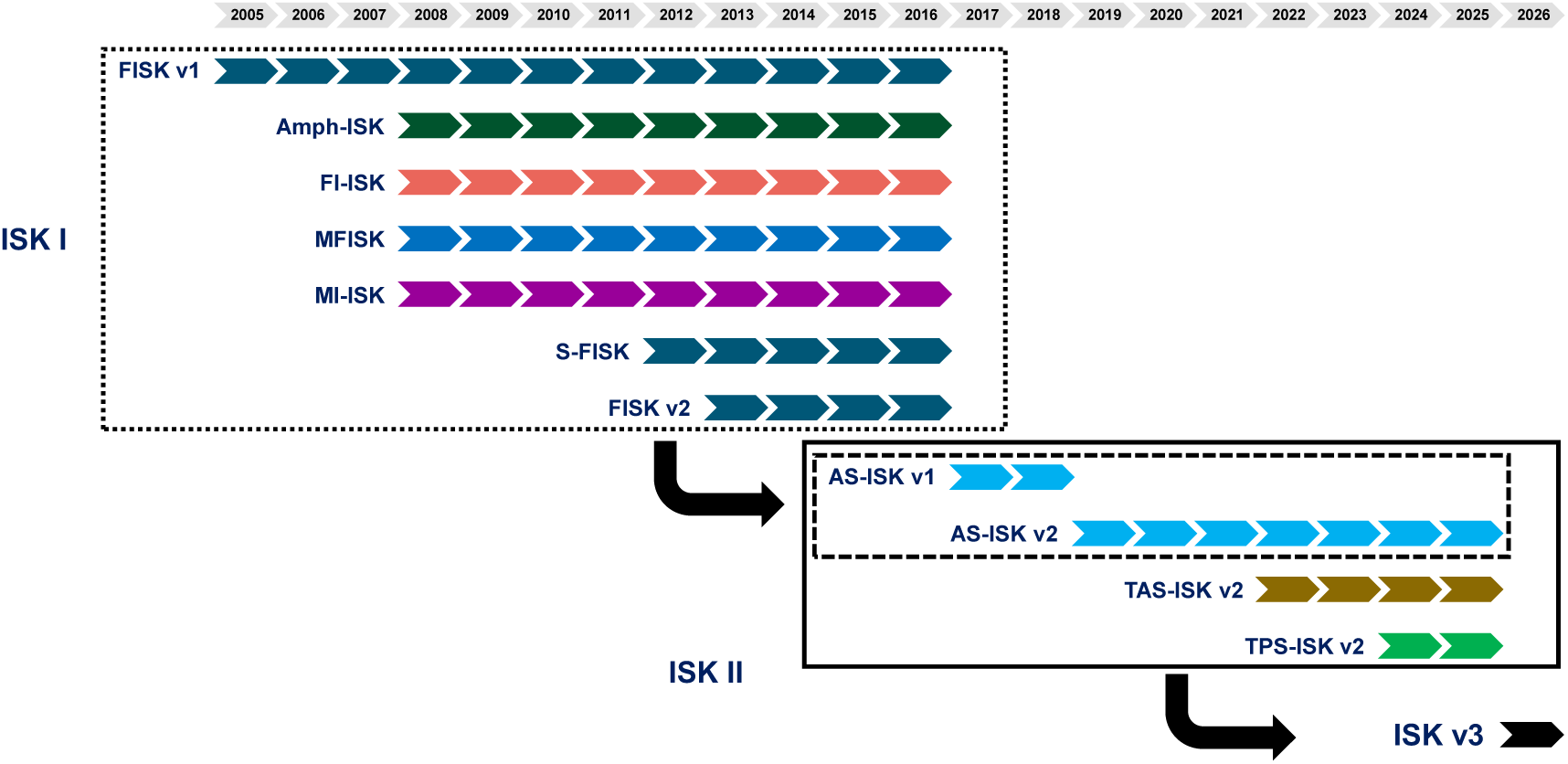
Evolution of the ISK family of risk-screening tools from 2005 to 2026. The timeline shows the progression from the first-generation ISK I toolkits, including FISK, Amph-ISK, FI-ISK, MFISK, MI-ISK, S-FISK and FISK v2, through the second-generation ISK II applications AS-ISK, TAS-ISK and TPS-ISK, culminating in the integrated ISK v3 platform.

Other non-native species risk-screening tools and frameworks have also been developed outside the ISK lineage. These include Harmonia+ (D’hondt et al. 2015), the Canadian Marine Invasive Screening Tool (CMIST: Drolet et al. 2016; Brown and Therriault 2022), the Fish Invasiveness Screening Test (FIST: Singh and Lakra 2011) and the more recently developed Non-Indigenous Species Screening Tool (NISST: Wilcox et al. 2025) and HeRA (Sapounidis et al. 2026). However, among available risk-screening protocols, the ISK framework is distinguished by its exceptionally broad documented application base, including 1,926 taxa screened from 209 applications across 266 risk assessment areas worldwide over the past 21 years (Vilizzi 2026a).

Because of their legacy, the second-generation tools AS-ISK, TAS-ISK and TPS-ISK extend the WRA-type approach across aquatic organisms, terrestrial animals and terrestrial plants. They share a common screening structure consisting of 55 questions divided into a Basic Risk Assessment (BRA) component of 49 questions, which are inherited from the first-generation ISK tools via the WRA, and an additional Climate Change Assessment (CCA) component of six questions. Upon completion of a screening, two scores are generated, namely the BRA and BRA+CCA, on the basis of which a species is ranked into low, medium or high risk for a defined risk assessment area. While the distinction between low and medium risk is based on the fixed threshold value of 1 (Pheloung et al. 1999), the distinction between medium and high risk is based on a threshold value obtained through calibration (Vilizzi et al. 2022a, 2022b). Finally, compared with the earlier self-automated Excel workbook architecture used by the WRA and first-generation ISK tools, and in contrast to the other risk-screening toolkits listed above, the second-generation ISK tools were developed as turnkey applications with a substantially improved graphical user interface (GUI), greater consistency of data entry and exchange, and multilingual support (Copp et al. 2021; Vilizzi et al. 2025a).

Despite their widespread use and global application (Vilizzi 2026a), AS-ISK, TAS-ISK and TPS-ISK were until now distributed and operated as separate software environments. This reflected their historical development, but created fragmentation in code maintenance, software distribution and user support. As the ISK framework expanded across organismal scopes, the need increased for a single integrated platform that would preserve the established questionnaires and scoring logic while improving consistency, transparency, deployment and usability, and strengthening reproducibility across the three toolkits.

ISK v3 was developed to address this need by integrating AS-ISK, TAS-ISK and TPS-ISK into a single platform. The aim was not to revise the scientific basis of the (multilingual) questionnaires, which recently underwent upgrade and alignment (Vilizzi et al. 2025a), but to consolidate the software environment around them. The ISK v3 platform therefore preserves the established screening structure of AS-ISK, TAS-ISK and TPS-ISK while introducing harmonized workflows for database creation, opening, conversion, screening management, taxonomic verification, threshold calibration and management, risk summaries, reporting, merging and export.

This paper describes the development, architecture and functionality of ISK v3, an integrated multilingual decision-support platform for non-native species risk identification. It highlights the software advances over the earlier second-generation ISK versions and explains how the platform supports harmonized risk identification workflows across aquatic organisms, terrestrial animals and terrestrial plants. Throughout, the term ‘toolkit/organismal scope’ refers to the broad organismal coverage of each of the three ISK toolkits: aquatic organisms in AS-ISK, terrestrial animals in TAS-ISK and terrestrial plants in TPS-ISK. By contrast, ‘organismal group category’ refers to the operational categories within each toolkit that are used for database organization, threshold management, risk summaries and reporting.

## Software design and development rationale

### General design principles

ISK v3 was designed around software integration, transparency and data-management principles. Its primary objective was to integrate AS-ISK, TAS-ISK and TPS-ISK into a single platform while preserving the questionnaire structure and toolkit-specific identity of the three established decision-support tools. Previous versions of the aquatic, terrestrial animal and terrestrial plant toolkits were distributed and operated as separate software environments (Copp et al. 2016, 2021; Vilizzi et al. 2022c, 2024). ISK v3 brings these three decision-support tools together within one entry platform. This facilitates code maintenance, software release and distribution, reduces fragmentation and provides users with a common workflow irrespective of organismal scope.

A central design requirement was to preserve the established ISK questionnaires and scoring logic. The questionnaire structure of AS-ISK, TAS-ISK and TPS-ISK was fully retained to ensure continuity with previous applications of the three toolkits. Existing users can therefore continue to apply the same toolkit-specific multilingual questionnaires (Vilizzi et al. 2025a), response structure, confidence scoring and risk-score interpretation (Vilizzi and Piria 2022; Vilizzi et al. 2022a), while benefiting from a redesigned and integrated software environment. Although AS-ISK, TAS-ISK and TPS-ISK are accessed through a single platform, each toolkit therefore retains its own database structure, organismal group categories and compatibility requirements. AS-ISK supports 27 organismal group categories, TAS-ISK supports 12 and TPS-ISK supports seven (Vilizzi et al. 2025a). Toolkit identity is further maintained through internal v3 database markers that allow the respective databases to be managed through a harmonized interface while preventing inappropriate use of a database within the wrong toolkit environment.

Another design goal was to consolidate and improve the multilingual interface displayed at runtime, including the prompts and messages shown while the software is being used, across the three toolkits. Multilingual implementation has been a core component in the development of the second-generation ISK tools (Copp et al. 2021; Vilizzi et al. 2025a, 2026b) and ISK v3 extends this functionality across its integrated platform. Runtime prompts and messages were revised to improve clarity and consistency across languages and toolkits.

ISK v3 was also designed to support threshold setting and calibration within the software, thereby integrating this main component of the risk identification process that previously required separate implementation outside the toolkits. Threshold handling was revised to remove the inclusion of the incorrect BRA+CCA threshold for risk-ranking of species under future climate conditions (see Vilizzi and Piria 2022) and was expanded so that users can manage thresholds by risk assessment area and organismal group category, apply available generalized thresholds (after Vilizzi et al. 2021) where appropriate and, whenever possible (Vilizzi et al. 2022a), conduct threshold calibration directly within the software. This links screening scores more explicitly to risk outcomes and reduces the need for external processing when selecting, calibrating or interpreting thresholds (Vilizzi et al. 2022a).

Database handling was significantly expanded to support more streamlined reporting, clearer presentation of risk outcomes, exchange and reuse of screening datasets, and collaborative workflows in which screening records produced by different assessors or databases can be merged after validation. As in the second-generation ISK tools, ISK v3 supports the creation and opening of toolkit-specific databases, the import and conversion of compatible v2 and AS-ISK v1 databases, noting that no v1 databases exist for TAS-ISK and TPS-ISK given their later development relative to AS-ISK, and the import of legacy aquatic first-generation ISK databases into AS-ISK. New features include merging of compatible screening records from separate databases and compact export of validated screening records and associated risk-outcome summaries. These functions were designed to reduce manual database manipulation and facilitate the consistency, traceability and reusability of screening outputs, which were previously limited in the second-generation ISK tools.

Unlike the second-generation ISK tools in which the password-protected database worksheets remained visible (though not editable) within the workbook structure, ISK v3 hides the underlying database sheets from the user interface. This separates internal data storage from user-facing data access and reduces the risk of accidental modification of protected database structures. Because users can now examine screening outputs through the Risk summary option and extract datasets through the Export function, they can work with the relevant data without interacting directly with the internal database worksheets.

ISK v3 also supports more flexible use of Excel while preserving controlled ISK workflows. As a turnkey application, it operates as a protected, dialog-driven application within its own Excel instance, but users can now open within ISK v3 separate Excel instances where needed, for example to view generated reports, risk summaries, exported databases or other supporting material. This improves usability without compromising database integrity, controlled data entry or the internal logic of the screening tools.

Taken together, these developments reflect both the planned integration of the ISK framework and the cumulative feedback received from users since the release of AS-ISK and, subsequently, TAS-ISK and TPS-ISK.

### Software environment

ISK v3 is implemented as a standalone application consisting of a Microsoft Excel macro-enabled workbook (.xlsm) with a Visual Basic for Applications (VBA)-based GUI. It was developed for the desktop version of Microsoft Excel 365 running on Windows 10 or Windows 11 and is intended to be saved and launched from a local folder on the user’s computer. This requirement reflects the reliance of the application on VBA procedures, Windows-compatible Excel controls and local workbook operations that are not consistently supported, or are disabled for security reasons, in Excel for macOS, Excel for the web or cloud-based and browser-based spreadsheet environments such as OneDrive, SharePoint, Teams, Google Drive or Dropbox. Accordingly, ISK v3 is not designed for use in these environments.

Because ISK v3 relies on VBA procedures and form-based (dialog-driven) interface controls, Excel macros and the required controls must be enabled in the local Excel installation. On opening, the program performs preliminary compatibility checks to confirm that it is being run in a supported local Excel 365 for Windows environment before proceeding to the dialog-driven GUI. Because of the multilingual support of the ISK framework, the GUI is initialized in the local Office/Excel display language where available, otherwise English is used.

### Platform organization

ISK v3 is organized around a single Start dialog that acts as the entry point to the integrated platform. From this dialog, users select the toolkit to be used, namely AS-ISK for aquatic organisms, TAS-ISK for terrestrial animals or TPS-ISK for terrestrial plants. The Start dialog also provides access to general interface settings, including language, colour scheme, background contrast and display mode, as well as to the User Guide and supporting documentation.

After toolkit selection, ISK v3 opens a toolkit-specific Console that provides the main operational pathway for the selected toolkit, including creation of new databases or opening of existing databases into the Main Screening Workspace dialog. Although the three toolkits are accessed through a common platform, their databases remain toolkit specific. This preserves the separate toolkit scope, organismal group category structure and compatibility requirements of AS-ISK, TAS-ISK and TPS-ISK, while allowing users to work within a harmonized interface. Toolkit identity is also maintained through internal version markers embedded in the databases. These markers allow ISK v3 to distinguish among compatible toolkit-specific databases, apply the appropriate opening and validation procedures, and prevent databases from being used with the wrong toolkit environment.

The Start and Console dialogs also incorporate dedicated front-end graphics that provide a visual identity for the integrated ISK v3 platform and help distinguish the three toolkit entry points. These interface elements support orientation within the software while preserving a consistent visual structure across AS-ISK, TAS-ISK and TPS-ISK. Across the three toolkits, the interface follows shared organizational principles. The same general workflow is used for database opening, screening creation and editing, question entry, threshold handling, risk summaries, reporting, merging and export. This common structure reduces the learning curve for users working with more than one toolkit, while preserving the scientific and operational differences among aquatic organisms, terrestrial animals and terrestrial plants.

## Main software components

### Start

ISK v3 is organized around a unified Start dialog (Fig. 2a) that provides the common entry point to the platform, allowing users to accept or select the interface language, colour scheme, background contrast, display mode and the required toolkit: AS-ISK for aquatic organisms, TAS-ISK for terrestrial animals or TPS-ISK for terrestrial plants. It also provides access to the program Overview, Credits, User Guide and dialog-specific Help file. The interface supports 31 languages (English, Albanian, Arabic, Bulgarian, simplified Chinese, Croatian, Czech, Dutch, Filipino, French, Georgian, German, Greek, Hebrew, Hungarian, Italian, Japanese, Korean, Macedonian, Persian, Polish, Portuguese, Romanian, Russian, Slovak, Slovenian, Spanish, Swedish, Thai, Turkish, Vietnamese), with the local Microsoft Office/Excel display language used by default where available, otherwise English. Display settings, including Colour scheme, Background contrast and Full or Compact view, affect the visual presentation and usability of the graphical interface.

**Figure 2.**
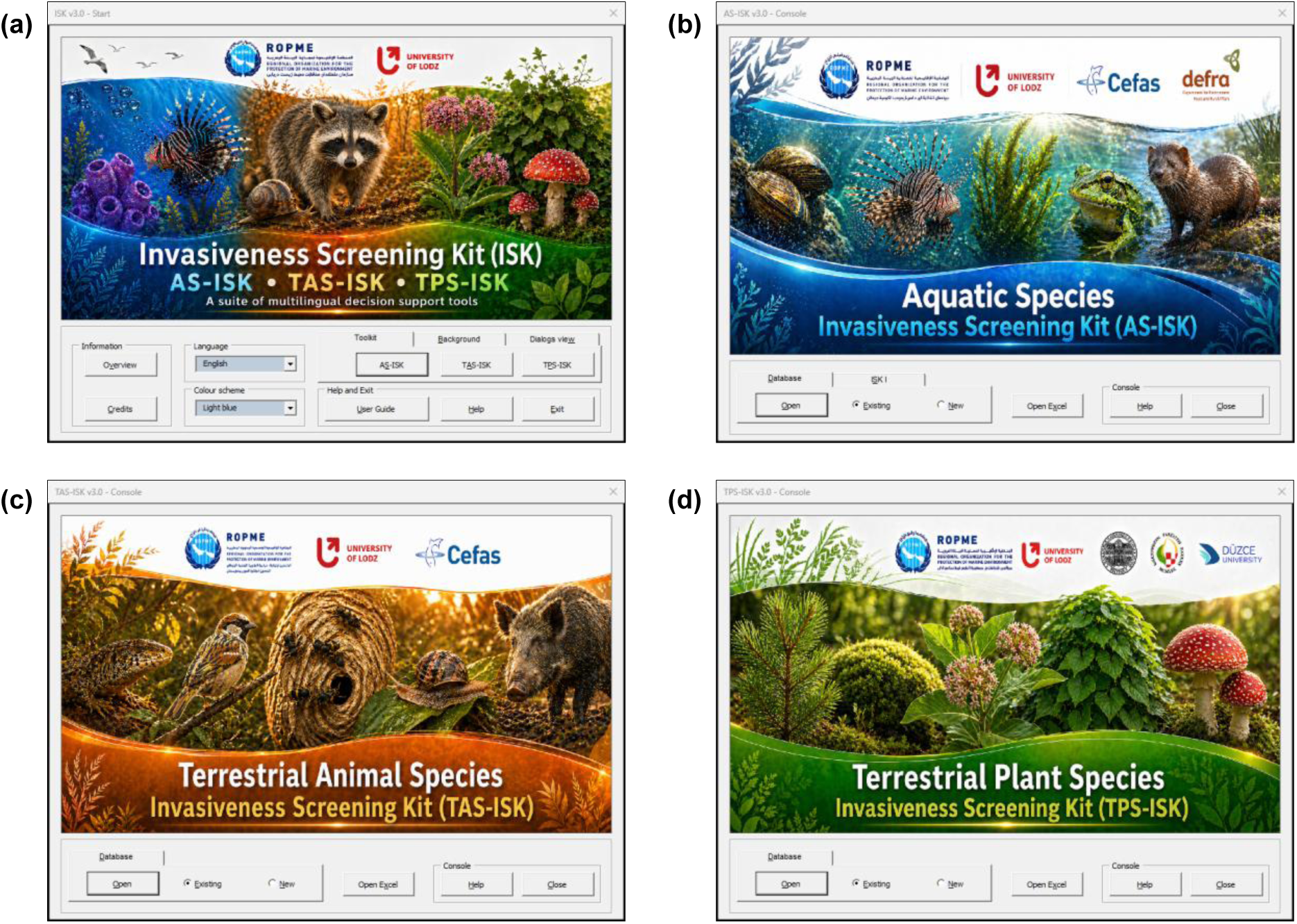
Entry dialogs of the Invasiveness Screening Kit (ISK) v3 platform. (a) Start dialog. (b) Aquatic Species Invasiveness Screening Kit (AS-ISK) Console. (c) Terrestrial Animal Species Invasiveness Screening Kit (TAS-ISK) Console. (d) Terrestrial Plant Species Invasiveness Screening Kit (TPS-ISK) Console.

### Consoles

Once a toolkit is selected, the corresponding Console dialog (Fig. 2b–d) is opened. The Console acts as the operational entry point for the selected toolkit, providing access to database creation, opening of existing compatible databases, integrated Excel access and launch of the Main Screening Workspace. The AS-ISK, TAS-ISK and TPS-ISK Consoles follow the same general organization, thereby providing a consistent workflow across toolkits, while retaining toolkit-specific database structures, organismal group categories, compatibility checks and opening procedures. For AS-ISK, the Console also includes an import option for legacy aquatic first-generation ISK databases. This allows previous screening databases from FISK v1, FISK v2, S-FISK (the Spanish version of FISK), Amph-ISK, FI-ISK, MFISK and MI-ISK to be imported into AS-ISK so that they can be updated to the current AS-ISK protocol. During database opening or import, ISK v3 performs preliminary formatting, consistency, duplicate-entry and re-scoring checks before screening records are loaded into the Main Screening Workspace. This provides continuity with previous first-generation ISK applications while bringing compatible historical records into the integrated v3 environment.

### Main Screening Workspace

The Main Screening Workspace dialog (Fig. 3) is the central environment in which screening records are managed after a database has been opened or created. It contains the Screening List and provides access to the main functional pages of ISK v3 (Screenings, Threshold, Risk summary, Report, Data tools) and to the Q&A dialog. The Screening List displays the records contained in the active toolkit database and includes both screening record metadata (i.e. risk assessment area, organismal group category ID, taxon name, common name, assessor) and calculated statistics and outputs (i.e. number of answered questions, threshold value, BRA and BRA+CCA scores and corresponding risk level outcomes, and date and time of creation or update).

**Figure 3.**
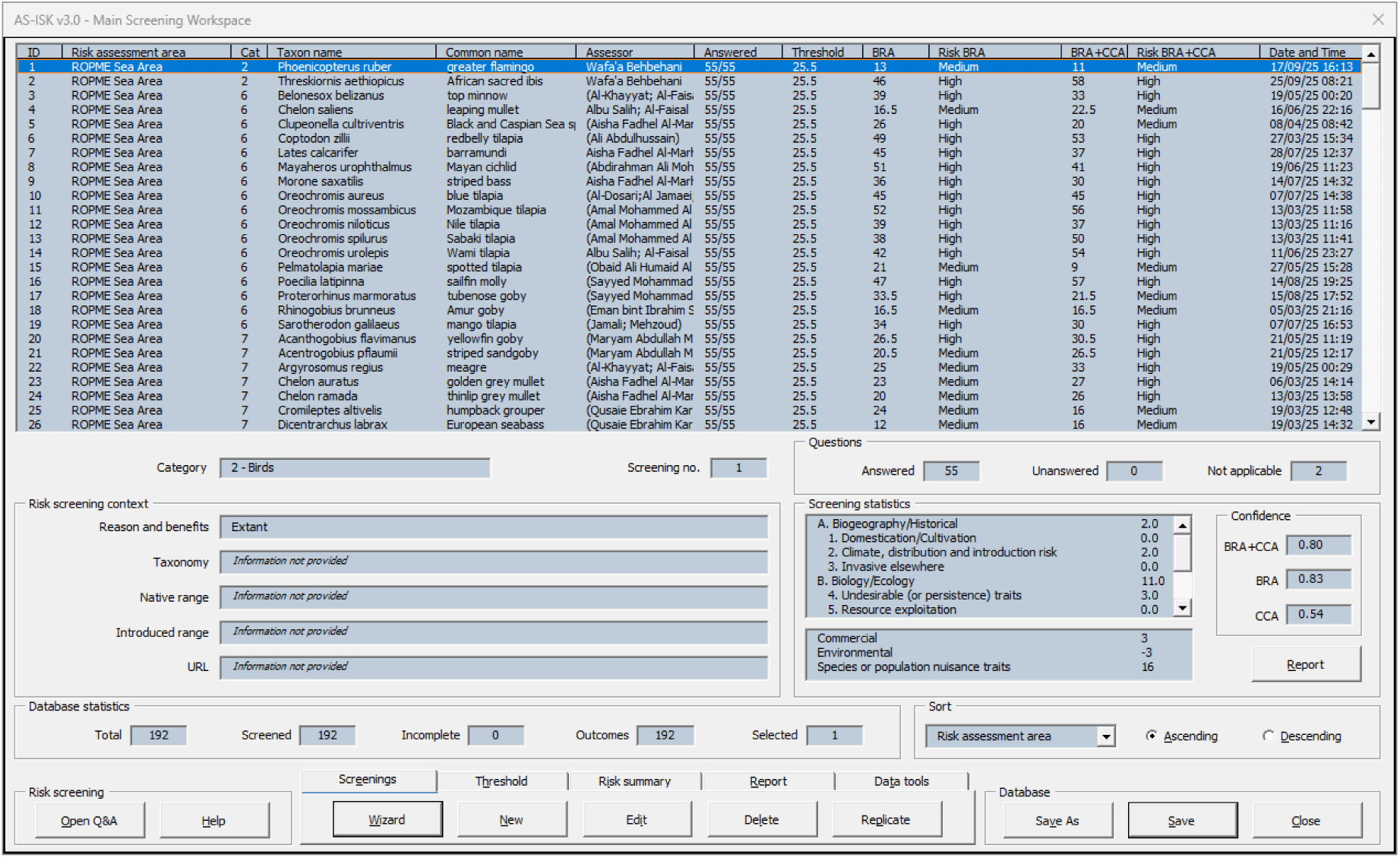
Main Screening Workspace of ISK v3, shown here for an AS-ISK database.

This dialog also provides contextual information for the selected screening record, including organismal group category description, screening number, question status (Answered, Unanswered, Not applicable), optional risk screening context (Reason and benefit, Taxonomy, Native range, Introduced range, URL), screening statistics (breakdown of scores across questionnaire sections and categories), confidence levels (for the BRA+CCA and separately for the BRA and the CCA) and database statistics. Sorting and selection controls support navigation through screening sets, while the tabbed structure separates record creation and editing, threshold management, risk summary generation, report production and database-level tools. This organization makes the Main Screening Workspace the main point of interaction between the user and the screening database. As in the second-generation ISK tools, each ISK v3 database supports up to 10,000 screening records. This upper limit was retained as a practical design capacity rather than as an expected database size, providing ample headroom for large multi-taxon, multi-assessor or multi-area applications while preserving compatibility with the workbook-based database architecture. The new risk summary, reporting, merge and export functions provide more practical ways to manage, review and reuse such outputs.

#### Screenings

This page provides the main tools for creating and managing screening records. Users can use the Wizard to generate multiple blank screening templates, create a new screening, edit an existing one, batch edit selected fields across multiple screenings, delete records or replicate selected fields from an existing record. These functions support both individual screening studies and larger multi-taxon (same or different organismal groups) or multi-assessor workflows also involving multiple risk assessment areas.

The Wizard dialog (Fig. 4) is used to create multiple screening templates with shared metadata, including organismal group category, risk assessment area and assessor. This is useful when several taxa for a certain organismal group are to be screened for the same risk assessment area. The New/Edit dialog (Fig. 5) is then used to enter or revise the core metadata for each screening record, including taxon name, common name, and, optionally, reason and socio-economic benefits, taxonomy, native range, introduced range and source URL.

**Figure 4.**
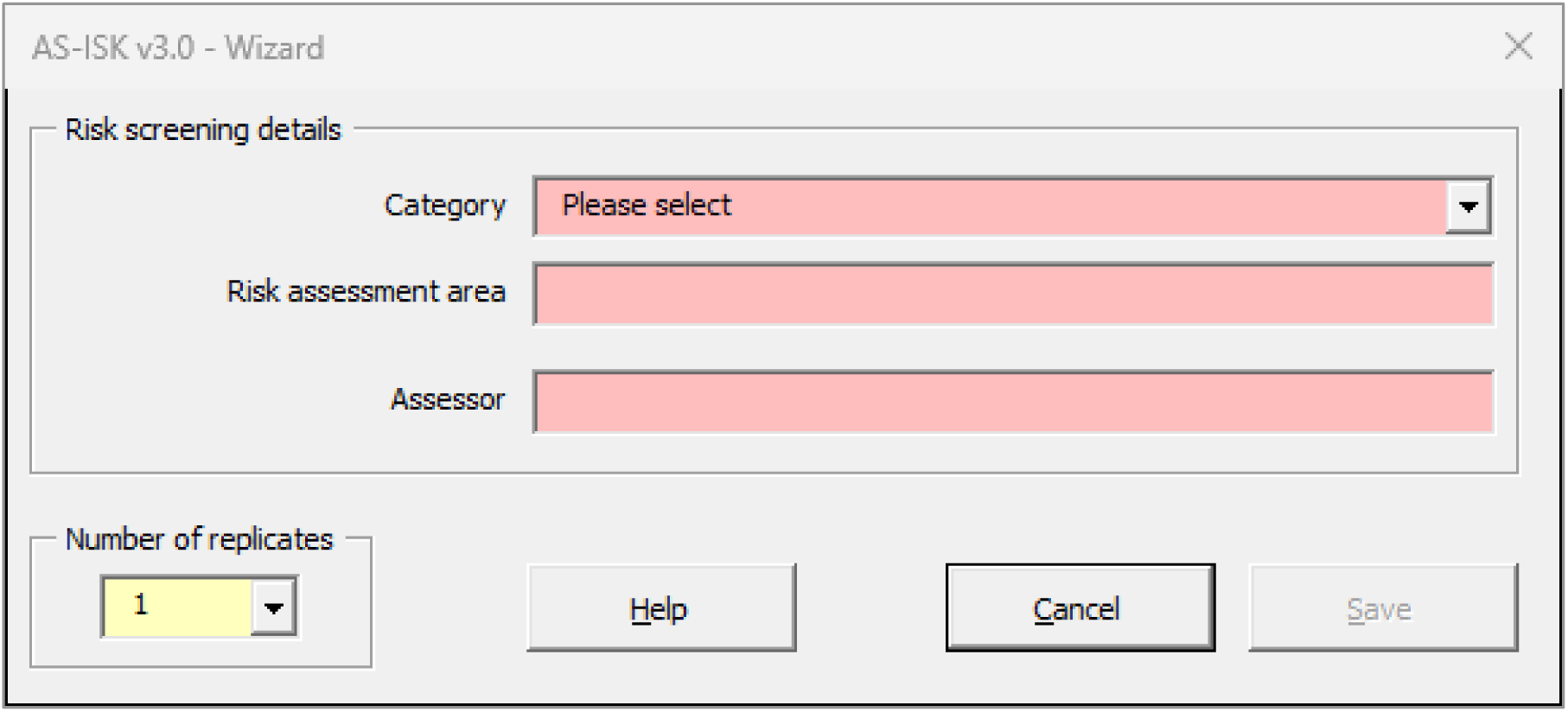
Wizard dialog of ISK v3.

**Figure 5.**
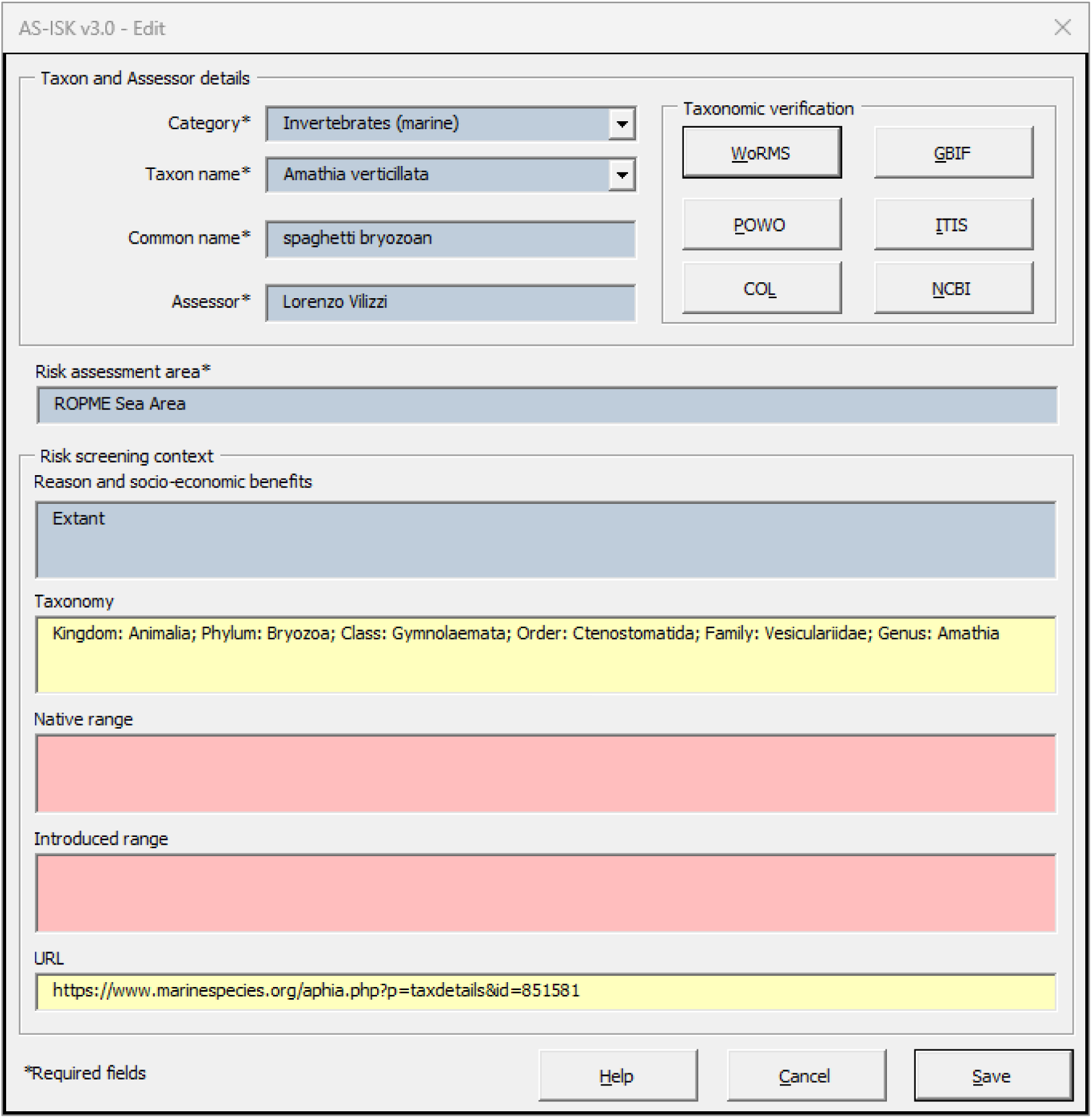
New/Edit dialog of ISK v3.

A new feature of the New/Edit dialog is the automatic compilation of 15,688 taxon names in the corresponding field based on the dataset (as of 22 May 2026) from the European Alien Species Information Network (EASIN: https://easin.jrc.ec.europa.eu/easin/). Taxonomic verification, another new feature of ISK v3, is integrated within the dialog. From this, users can verify taxon names through external taxonomic and biodiversity resources, including the World Register of Marine Species (WoRMS: https://www.marinespecies.org/), Global Biodiversity Information Facility (GBIF: https://www.gbif.org/), Plants of the World Online (POWO: https://powo.science.kew.org/), Integrated Taxonomic Information System (ITIS: https://www.itis.gov/), Catalogue of Life (https://www.catalogueoflife.org/) and National Center for Biotechnology Information Taxonomy (NCBI: https://www.ncbi.nlm.nih.gov/Taxonomy/Browser/wwwtax.cgi). Where available, accepted names and source information can be retrieved and used to update the screening record. This improves traceability of taxonomic decisions and reduces the risk of inconsistent or outdated names. However, the function is intended as decision support rather than as an automatic replacement for taxonomic judgement: the user decides whether suggested updates are applied.

The Replicate dialog (Fig. 6) allows users to generate one or more new screening records from a screening selected in the Screening List. This option is useful when initiating screenings that share part of the same metadata. The organismal group category is always retained in the replicated screening, whereas the user can select which of the remaining fields are copied. Required fields that are not replicated are left to be completed in the new record. This provides a controlled way to create related screening records while reducing repetitive data entry and limiting the risk of accidental duplication.

**Figure 6.**
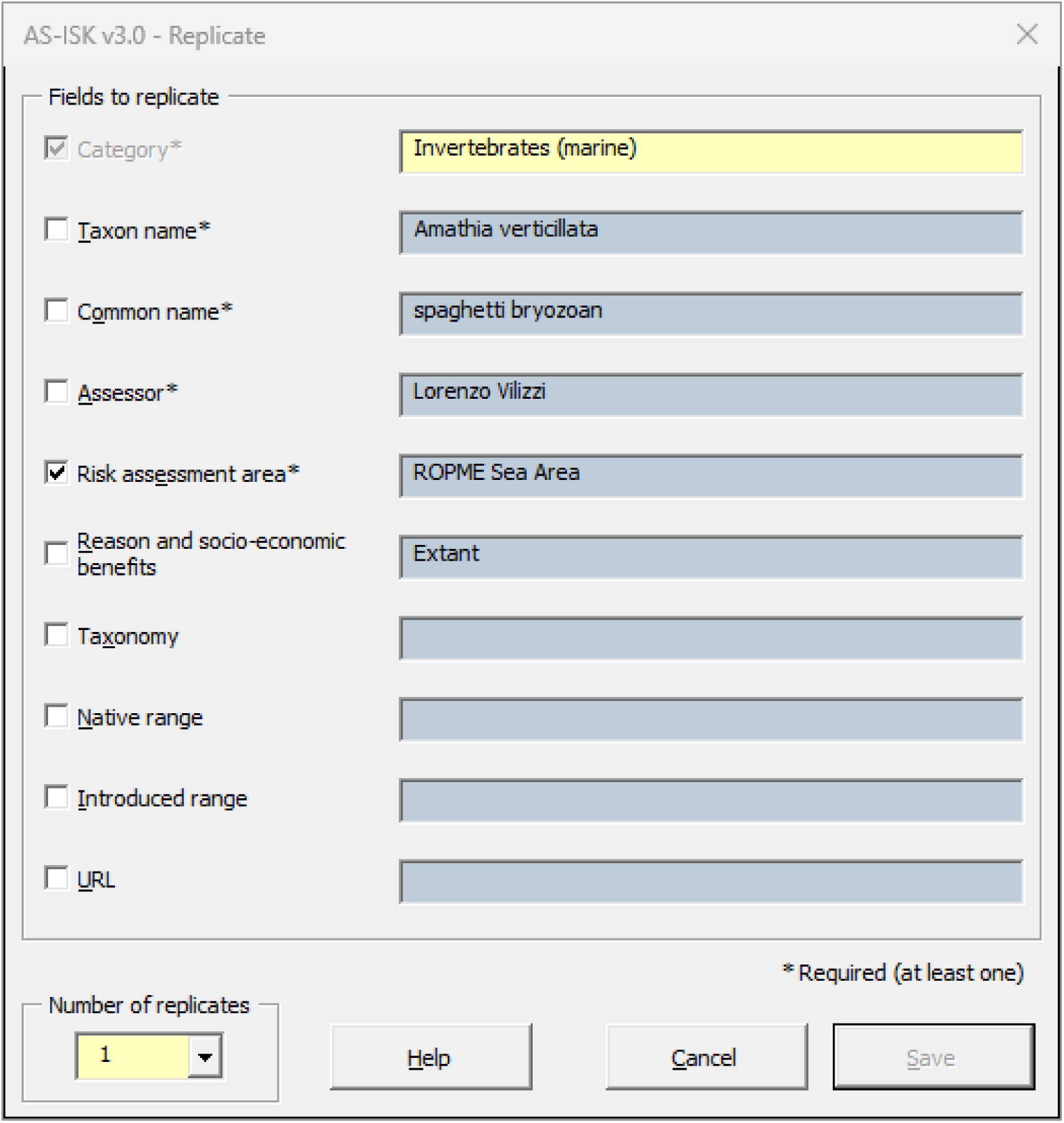
Replicate dialog of ISK v3.

#### Threshold

This page supports the interpretation of screening scores by allowing thresholds to be assigned and managed by risk assessment area and organismal group category. Users can set, reset or clear thresholds, apply available generalized thresholds where appropriate and use thresholds to interpret both BRA and BRA+CCA scores. Where multiple screenings are available for the same taxon in the same risk assessment area, ISK v3 supports interpretation through averaged scores and corresponding outcomes. A major addition in ISK v3 is the integration of Receiver Operating Characteristic (ROC) curve-based threshold calibration directly within the software (see Vilizzi et al. 2022a). This allows users to calibrate screening thresholds using built-in *a priori* invasive and non-invasive categorizations (after Vilizzi 2026a) without having to export the data to external statistical software. The ROC workflow produces calibration statistics including the area under the curve, 95% confidence intervals, sensitivity, specificity, Youden’s *J* and confusion matrix outputs. Where required, threshold summary outputs and ROC-related reports can also be generated. This integration links threshold setting more directly to the global *a priori* categorization dataset recently developed to support calibration of invasion risk screening applications (Vilizzi 2026a). By embedding ROC computation in the same platform used for database management and screening, ISK v3 makes threshold calibration more accessible, transparent and reproducible, particularly for users managing larger screening datasets across multiple taxa, organismal group categories or risk assessment areas.

#### Risk summary

This page provides synthesis outputs for screening datasets. Summaries can be generated by risk assessment area, taxon or organismal group category, and include BRA and BRA+CCA scores and, if calibrated, corresponding risk outcomes. Where multiple screenings exist for the same taxon within the same risk assessment area, summary outputs use averaged scores to support interpretation across repeated screenings from different assessors. These outputs provide a structured basis for comparing taxa, areas and organismal group categories, and help users move from individual screening records to dataset-level interpretation.

#### Report

This page generates structured documentation for screening records. Reports can be produced as single, separate or grouped outputs, with file type selection available according to user requirements. This function supports communication, archiving, review and sharing of screening results, while preserving the underlying structure of the screening record, including metadata, scores, outcomes and supporting information.

#### Data tools

This page contains database-level management functions. These include merging compatible databases, exporting compact database outputs, re-scoring the active database and opening a separate Excel instance when other workbooks need to be consulted outside the controlled ISK v3 session. The Merge function supports collaborative and multi-assessor workflows by allowing up to 50 compatible ISK v2 and v3 databases to be combined after validation checks for formatting, compatibility, duplicate entries, mandatory fields and internal consistency. The Export function produces cleaned database outputs that retain key screening fields and calculated results while removing internal working structures. This makes the exported files more suitable for archiving, downstream analysis, data sharing and integration into wider screening or reporting workflows.

#### Q&A

This dialog (Fig. 7) is used to enter responses to the 55 ISK questions, comprising the BRA and CCA components. For each question, the user records the response, confidence level, scope and justification, with navigation controls allowing movement through individual questions or direct access to a selected question. The interface tracks answered, unanswered and not applicable questions, and screening statistics are updated accordingly. The questionnaires and scoring logic have not been changed in ISK v3, ensuring continuity with previous AS-ISK, TAS-ISK and TPS-ISK applications. A new feature is the indication for each question of its Scope, i.e. whether it refers to the taxon’s characteristics in general or to its introduced range or specifically to the risk assessment area under investigation.

**Figure 7.**
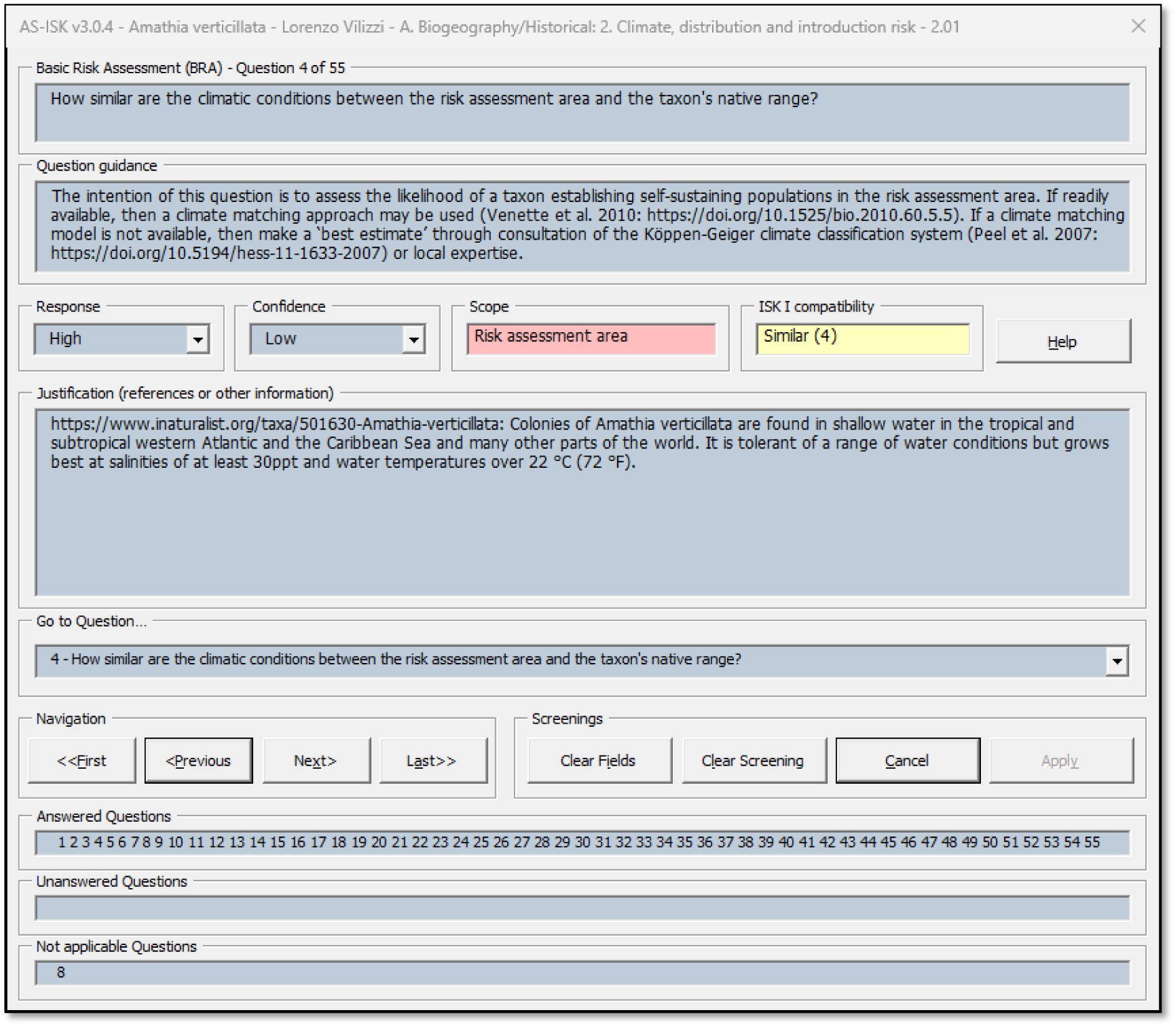
Q&A dialog of ISK v3.

### Workflow example

A typical ISK v3 workflow begins when the user opens the macro-enabled workbook in a supported local Microsoft Excel 365 desktop environment under Windows 10 or Windows 11. After the preliminary compatibility checks, the Start dialog is displayed. From this entry point, the user accepts (or selects) the interface language, colour scheme, background contrast and display mode, and then chooses the appropriate toolkit: AS-ISK for aquatic organisms, TAS-ISK for terrestrial animals or TPS-ISK for terrestrial plants.

The selected toolkit opens its corresponding Console, from which the user either creates a new database or opens an existing compatible database. In AS-ISK, this step may also include importing a legacy aquatic first-generation ISK database so that previous screenings can be updated to the current AS-ISK protocol. Once the database has been created, opened or imported, ISK v3 launches the Main Screening Workspace and displays the active Screening List.

In the Main Screening Workspace, screening records can then be added individually, edited from existing records, replicated or generated as multiple templates through the Wizard. For each screening, the user enters the main mandatory metadata, including the organismal group category, taxon name, common name, assessor and risk assessment area, and optionally the reason for screening, taxonomy, native range, introduced range and source URL. Where required, the user can verify taxon names through online taxonomic and biodiversity resources and decide whether to apply any suggested update to the screening record, including taxonomy and source URL information.

The user then completes the Q&A workflow by answering the 55 ISK questions, recording the response, confidence level, scope and justification for each question. The software tracks answered, unanswered and not applicable questions and updates the corresponding screening statistics and risk scores. After the screening records have been completed, the user can assign existing thresholds, apply available generalized thresholds or perform ROC curve-based calibration to derive thresholds from the available *a priori* invasive and non-invasive categorizations.

Finally, ISK v3 supports interpretation and output generation through the Risk summary, Report and Data tools pages. The user can generate summaries by risk assessment area, taxon or organismal group category, produce structured reports for screening records and export compact cleaned datasets retaining key metadata and calculated outputs. This workflow links data entry, evidence documentation, taxonomic checking, scoring, threshold interpretation, reporting and export within a single integrated software environment.

The workflow can be tested using the supplementary materials provided with this article: sample native ISK v3 databases prepared from published ISK application datasets (Suppl. materials 1–3), original second-generation ISK databases for testing conversion into ISK v3 (Suppl. materials 4–6), a legacy FISK v2 database for testing import into AS-ISK v3 (Suppl. material 7), TPS-ISK v2 databases for testing the Merge function (Suppl. materials 8–12), example outputs generated by ISK v3 (Suppl. materials 13–18) and a reviewer guide with source-publication information and suggested test workflows (Suppl. material 19).

## Application relevance and use cases

### Research applications

ISK v3 supports a range of research applications in invasion science and applied biosecurity by providing a harmonized platform for risk screening across aquatic organisms, terrestrial animals and terrestrial plants. It can be used in screening studies involving single species, multiple species from the same organismal group or larger screening datasets spanning different organismal groups, ecosystems, pathways, regions or risk assessment areas. Previous ISK applications have included freshwater, brackish and marine taxa; taxon-specific and taxon-generic screening exercises; regional species-prioritization studies; comparisons among risk assessment areas; pathway- or sector-focused applications involving aquaculture, ornamental trade, pet trade, ballast water and other introduction routes; and, in the case of the second-generation ISK tools, studies incorporating current and future climate conditions (for a full list of applications, see Vilizzi 2026a). Because the three toolkits are accessed through a common software environment while retaining their established questionnaires and scoring logic, ISK v3 facilitates more consistent application of WRA-type screening principles across toolkit scopes and provides a common framework for comparative risk screening, calibration studies and synthesis or meta-analysis of screening outputs (Vilizzi et al. 2019, 2021; Hill et al. 2025; Vilizzi 2026a).

The platform is also relevant to calibration studies (Vilizzi et al. 2022a). The integration of ROC curve-based threshold calibration within the software allows researchers to estimate, compare and document thresholds without relying on separate statistical workflows. This is particularly useful where screening outcomes need to be evaluated against *a priori* invasive or non-invasive categorizations, or where calibrated thresholds are compared among risk assessment areas or organismal groups.

The database, report and export functions also support synthesis of screening outputs. Screening records can now be managed consistently, exported in compact form and used in downstream analyses, including comparative studies and meta-analyses of screening results. In this way, ISK v3 provides a common framework for generating, organising and reusing screening information, while improving traceability of the evidence, assumptions and justifications underlying individual screening decisions.

### Management and policy applications

ISK v3 is equally relevant to applied biosecurity workflows, where rapid, transparent and repeatable risk identification is needed to support decision-making (Canavan et al. 2025). The tool can assist early warning and horizon scanning by helping users identify non-native species that may require closer attention before or after introduction (Nunes et al. 2025). It can also support prioritization for monitoring, surveillance or follow-up risk assessment, especially when management resources are limited and many taxa must be considered (e.g. O’Shaughnessy et al. 2023; Glamuzina et al. 2024; Vilizzi et al. 2026a).

Because each screening record includes structured metadata, responses, confidence scores, scope information and written justifications, ISK v3 provides a documented basis for explaining how screening outcomes were reached (Vilizzi and Piria 2022). This is important when results are used to inform management or policy discussions, where transparency and reproducibility are essential, including the provision of reports in languages other than English (Copp et al. 2021). The Risk summary and Report functions further support communication by converting screening records into structured outputs that can be reviewed by technical experts, managers or decision makers.

At regional scale, ISK v3 can contribute to biosecurity planning by providing a consistent workflow for compiling and updating screening records across taxa, risk assessment areas and organismal group categories (e.g. Vilizzi et al. 2026a). This allows risk-screening information to be compared and summarized in a way that supports prioritization, monitoring design and the identification of species requiring additional evidence or management attention.

### Country-level adoption and institutional applications

ISK v3 is not restricted to any single region, country or institution. Its structure is intended to support research groups, government agencies, regional organizations and technical working groups that require a shared decision-support framework for non-native species risk identification. Institutions can use the platform to apply common screening procedures, harmonize documentation among assessors, generate comparable outputs and maintain screening databases that can be updated as new evidence becomes available.

A country-level example of institutional uptake is provided by the Philippines, where AS-ISK has been adopted as an official decision-support tool by the Bureau of Fisheries and Aquatic Resources and implemented by the National Fisheries Research and Development Institute for screening introduced aquatic species (Republic of the Philippines 2021). Although this adoption concerns AS-ISK rather than ISK v3 specifically, it illustrates how ISK-based workflows can be embedded within national biosecurity and fisheries-management systems. Such applications require not only a scientifically validated screening protocol, but also a practical software environment capable of supporting assessor training, standardized documentation, database maintenance and communication of screening outcomes.

ISK v3 extends this institutional potential by providing a single integrated platform for aquatic organisms, terrestrial animals and terrestrial plants. This makes it suitable for programmes that need to apply comparable risk-identification workflows across multiple organismal groups, risk assessment areas or administrative jurisdictions. The platform can therefore support national or regional screening programmes in which multiple assessors contribute records to shared databases, thresholds are applied consistently and outputs are suitable for technical review, reporting and policy discussion.

At regional scale, ISK v3 is relevant to biosecurity contexts in which comparable screening procedures are required across jurisdictions. The Regional Organization for the Protection of the Marine Environment (ROPME: ropme.org) Sea Area provides one such example, as coordinated risk-identification workflows are needed across Member States for marine non-native species occurring, spreading or potentially arriving within a shared regional sea (Vilizzi et al. 2026a). More generally, the integration of AS-ISK, TAS-ISK and TPS-ISK into a single platform provides practical support for regional or institutional programmes that require standardized data-entry structures, taxonomic verification, threshold handling, risk summaries, reporting functions and exportable screening datasets.

This institutional relevance illustrates one of the main intended uses of ISK v3: coordinated biosecurity initiatives in which standardized, transparent and well-documented screening workflows are needed across multiple taxa, assessors and risk assessment areas. By combining toolkit integration, database management, taxonomic verification, threshold handling, reporting and export functions, ISK v3 provides a practical software infrastructure for institutional adoption and harmonized implementation of non-native species risk-identification procedures.

### Availability, versioning and citation

ISK v3 is archived on Zenodo, where software packages are deposited as versioned releases. The recommended software citation uses the Zenodo concept DOI, which resolves to the latest release in the ISK v3 software line (Vilizzi 2026b). This approach allows users to cite the current ISK v3 platform consistently, while preserving access to individual versioned releases for reproducibility, documentation and software-history purposes. The associated Zenodo record includes the macro-enabled workbook, User Guide, Help files, changelog and citation information. The User Guide provides installation requirements, interface descriptions and operational instructions, while the Help files provide context-specific guidance for the Start dialog, toolkit Consoles, Main Screening Workspace and associated data-entry dialogs. The supplementary materials further support software review and reproducibility by providing sample databases, conversion and import examples, merge test files, representative ISK v3 outputs and a reviewer guide.

In addition to citing the software, users are encouraged to cite the relevant methodological sources according to the toolkit used and the context of application. AS-ISK applications should cite Copp et al. (2016, 2021), Vilizzi and Piria (2022), and Vilizzi et al. (2021, 2025a); TAS-ISK applications should cite Vilizzi and Piria (2022) and Vilizzi et al. (2022c, 2025a); and TPS-ISK applications should cite Vilizzi and Piria (2022) and Vilizzi et al. (2024, 2025a). Where ROC curve-based threshold calibration is used, the relevant calibration protocol should also be cited (Vilizzi et al. 2022b). When the built-in ROC calibration workflow is used with the global *a priori* categorization dataset, this dataset should be cited separately (Vilizzi 2026a). This citation structure distinguishes among the software platform, the methodological basis of the individual toolkits, the calibration protocol and the dataset used for threshold calibration, thereby supporting both software attribution and methodological traceability.

## Discussion

### Main contribution

ISK v3 represents the latest step in the evolution of the ISK tools from a set of related but separate multilingual decision-support tools into a single integrated platform. Previous versions of AS-ISK, TAS-ISK and TPS-ISK were developed sequentially to cover all major organismal scopes and were distributed as separate software environments. ISK v3 consolidates these tools within one platform while preserving their scoring logic, established questionnaires and toolkit-specific scopes (Vilizzi et al. 2022a, 2025a, 2026b). Overall, this follows the same principle that led to the merging of the first-generation aquatic ISK tools into AS-ISK (Copp et al. 2016), which allowed users to benefit from the advantages of a single platform while retaining broad applicability across aquatic organismal groups. ISK v3 extends this consolidation principle further by bringing AS-ISK, TAS-ISK and TPS-ISK into a common software environment for aquatic organisms, terrestrial animals and terrestrial plants. This integration improves cross-toolkit consistency by providing a common entry point, shared GUI components and improved, streamlined data-analysis and data-handling capabilities. At the same time, the platform retains the separate identity of each toolkit through toolkit-specific databases, organismal group categories, validation checks and internal version markers. This balance between integration and toolkit specificity is central to the design of ISK v3.

A further contribution of ISK v3 is the streamlining and stabilization of the underlying codebase. The software was revised to improve consistency across AS-ISK, TAS-ISK and TPS-ISK, strengthen error handling and improve control of database operations. These changes are consistent with general principles of scientific-software development, including maintainability, robustness, transparency and error control (Wilson et al. 2014; Taschuk and Wilson 2017). The platform also preserves the reproducible structure of ISK screenings while strengthening transparency and traceability. Database validation, taxonomic verification, threshold management, ROC calibration, report generation and compact export functions support traceable workflows from data entry to interpretation and communication of results. By combining these functions in a multilingual software environment, ISK v3 makes the application of WRA-type screening principles more consistent, accessible and reusable across different users, regions and organismal groups.

### Advances over previous versions

ISK v3 introduces several advances over the previous standalone AS-ISK, TAS-ISK and TPS-ISK software environments. The most visible change is the integration of the three toolkits into one platform with a unified Start dialog, toolkit-specific Consoles and a common Main Screening Workspace. This reduces fragmentation in software distribution and use, while allowing users to move between toolkits through a shared interface structure.

A less visible but important advance is the substantial streamlining and stabilization of the underlying VBA code and the harmonization of the underlying spreadsheets. Together, these components underpin the GUI architecture of the second-generation ISK toolkits, which consists of tightly controlled dialogs forming the user interface, protected worksheets acting as the data-storage layer and code-based business logic mediating between the two (Copp et al. 2016). Large parts of the previous toolkit-specific code and underlying spreadsheets were revised, adapted or reorganized within the integrated v3 architecture, while preserving the established questionnaires, scoring logic and toolkit-specific database structures. The use of modal dialogs for key workflows, including the Wizard, New/Edit, Replicate and Q&A dialogs, further strengthens controlled data entry by requiring users to complete, save or cancel each operation before returning to the Main Screening Workspace. This revision improved consistency across AS-ISK, TAS-ISK and TPS-ISK, reduced duplication, strengthened error handling and enhanced control of database operations. It also made the software more stable, more maintainable and better able to manage incompatible files, incomplete operations and differences among user Excel installations without compromising the active database, in line with general principles for robust scientific software (Wilson et al. 2014; Taschuk and Wilson 2017).

A further advance is the integration of live online taxonomic verification within the screening-entry workflow. Previous ISK versions relied on user-entered taxon names and static database content, whereas ISK v3 allows users to query major internet-based taxonomic and biodiversity resources directly from the enhanced New/Edit dialog. This enables taxon names to be checked against external resources, accepted names to be retrieved where available and taxonomic source information to be documented in the screening preamble. By linking the screening workflow to live taxonomic resources, ISK v3 improves name standardization, taxonomic traceability and consistency of screening records, which are recognized challenges in biodiversity data management and taxon-name harmonization (Costello et al. 2013; Grenié et al. 2023). The feature remains user-controlled, since suggested updates must be confirmed by the assessor and are intended to support, rather than replace, taxonomic judgement.

The ISK v3 platform also improves database workflows by providing additional procedures for summarizing, merging and exporting screening databases, and expands the analytical and reporting capacity of the software. Built-in ROC curve-based threshold calibration allows users to estimate and document thresholds directly within the platform. Risk summary outputs support synthesis by risk assessment area, taxon and organismal group category, while reporting functions produce structured documentation of screening records. Merge and compact export tools support collaborative screening, downstream analysis, archiving and data sharing. Maintenance releases have also improved compatibility, interface behaviour, language handling and documentation, while leaving database structure and scoring logic unchanged.

### Limitations

ISK v3 has defined software-environment requirements. It is implemented as a Microsoft Excel macro-enabled workbook and is designed for the desktop version of Microsoft Excel 365 running on Windows 10 or Windows 11. It is not intended for Excel for Mac, Excel for the web or cloud-only spreadsheet environments. Users must therefore run the software from a local folder in a supported Windows Excel environment, with macros and required form controls enabled. These requirements reflect the use of VBA procedures, form-based interface controls and controlled database-handling workflows. They also allow ISK v3 to operate as an integrated dialog-driven screening platform rather than as a conventional spreadsheet. However, they mean that use in unsupported environments may prevent the software from opening or functioning as intended.

Some functions also depend on external conditions or user judgement. Online taxonomic verification requires internet access and availability of the relevant external taxonomic resources. It can assist with checking names and retrieving source information, but it does not replace taxonomic expertise or responsibility for the final taxonomic decision. Similarly, ROC-based threshold calibration depends on the quality, relevance and representativeness of the *a priori* invasive and non-invasive categorization data used. Calibration outputs should therefore be interpreted in relation to the dataset, taxa, risk assessment area and categorization criteria on which they are based.

### Future development

Future development of ISK v3 is expected to focus on maintenance, taxonomic support, threshold resources, multilingual expansion, continued usability improvements and controlled response-support functions.

Maintenance will remain a key component of future releases. This may include refinements to interface behaviour, additional compatibility adjustments for managed institutional Excel installations, and correction of language or terminology issues also in response to user feedback. Such maintenance releases would continue to improve stability and usability without altering the scientific structure of the toolkits or the interpretation of screening scores.

Taxonomic support is a second area for further development. ISK v3 already integrates online taxonomic verification through major taxonomic and biodiversity resources and includes automatic name completion based on the EASIN dataset available at the time of release. Future releases could expand this functionality beyond the current EASIN-based taxon-name resource by increasing the number and coverage of pre-compiled taxon names, improving matching of synonyms and accepted names, refining links to external taxonomic resources and incorporating additional curated datasets where appropriate. This would further improve taxonomic traceability and reduce the risk of inconsistent or outdated names, while retaining the principle that taxonomic verification supports, but does not replace, expert judgement.

Threshold resources could also be expanded. ISK v3 already allows users to assign, manage and calibrate thresholds by risk assessment area and organismal group category, and includes ROC curve-based calibration within the software. A future development would be to incorporate, where appropriate, thresholds already calibrated for specific risk assessment areas from previous applications of the ISK tools (see Vilizzi et al. 2022a). Such a curated threshold resource would need to retain clear provenance, including toolkit, organismal group, risk assessment area, calibration dataset, *a priori* categorization criteria and source reference. This would support users who wish to apply existing calibrated thresholds transparently, while preserving the option to recalibrate thresholds when new data, different taxa or different risk assessment areas require it.

Further multilingual expansion is another major future direction. ISK v3 currently supports 31 languages, including English, but a recently developed global framework for communication of biological invasion risks (Vilizzi et al. 2026b) provides a basis for expanding terminology and interface support to additional non-English languages where sufficient validation and expert review are available. Such expansion would be particularly relevant for countries and regions where biological invasion terminology is still developing, and where local-language decision-support tools may improve communication among scientists, competent authorities, managers and stakeholders. Future language updates may therefore include additional translations, terminology refinements, improved consistency of prompts and messages, and wider validation of key risk-analysis and decision-support terms.

Continued usability improvements would further support training, uptake and implementation. Additional examples, tutorials, worked workflows and dedicated ISK training workshops would be useful for users applying ISK v3 in regional screening programmes, institutional settings or comparative research studies. Previous main author-led training activities on ISK-based risk screening, including documented online and regional workshops for China, the ROPME Sea Area and Armenia (Beijing Lingyunyi Data Technology Co. 2024; AlMulla 2025; Soghoyan 2025), have illustrated the value of combining conceptual background, practical software use, data management and regionally relevant case studies for building capacity among assessors. Future workshops could further standardize practical use of the software, including database creation and management, screening entry, taxonomic verification, threshold calibration, interpretation of repeated screenings, merging of multi-assessor databases, preparation of compact exported datasets and production of reports for review or policy discussion.

Controlled response-support functions may also be developed. The inclusion of Scope in the Q&A workflow creates opportunities for future controlled support of pre-compiled response material for question components that are generic to a taxon or to its introduced range, while retaining assessor responsibility for risk assessment area-specific interpretation. This development would help streamline and accelerate the risk-screening process, particularly in large multi-taxon or multi-assessor applications where completion of the full ISK questionnaire can be time-consuming. Such functionality could reduce repetitive evidence gathering, improve consistency among screenings and increase user uptake, while preserving assessor judgement, documentation of evidence and final responsibility for each screening response. This direction would require careful curation and validation and should be developed without compromising the evidence-based and context-specific nature of ISK screenings.

## Conclusion

ISK v3 provides an integrated, multilingual and calibration-enabled platform for non-native species risk screening across aquatic organisms, terrestrial animals and terrestrial plants. By harmonizing workflows and improving database management, taxonomic verification, threshold calibration, summary generation, reporting and export functions, it strengthens the transparency, traceability and practical applicability of ISK-based risk identification for research, management and policy. The platform therefore represents both a consolidation of the second-generation ISK toolkits and a software foundation for their continued application, maintenance and development.

## Additional information

### Conflict of interest

The authors have declared that no competing interests exist.

### Ethical statement

No ethical approval was required because this article describes software development and did not involve human participants, animal experimentation or new field sampling.

### Artificial Intelligence (AI) use

OpenAI’s ChatGPT was used only for language editing, including grammar, spelling and wording improvements. Responsibility for the manuscript content rests entirely with the author.

### Funding

This work was funded by the Regional Organization for the Protection of the Marine Environment (ROPME), Kuwait, under Phase II (2026) of the 2025–2030 ROPME Strategic Programme “Risk Analysis of Non-native Species in the ROPME Sea Area”. The work corresponds to Deliverable D1 of the contractual framework, covering the development, beta testing and release of the ISK software. Funding supported the development, integration, testing, documentation and operational release of ISK v3.

### Authors contributions

L.V.: Conceptualization, Methodology, Software, Validation, Formal analysis, Visualization, Writing – original draft, Writing – review and editing. A.A-M.: Validation, Writing – review and editing, Project administration, Funding acquisition.

### Data availability

The ISK v3 software package, including the macro-enabled workbook, User Guide, Help files and changelog, is available from Zenodo through the ISK v3 concept DOI: https://doi.org/10.5281/zenodo.20414051. The global *a priori* categorization dataset used to support ROC curve-based threshold calibration is available separately from Zenodo: https://doi.org/10.5281/zenodo.19240779. Supplementary materials are provided to support software testing and reproducibility. Sample native ISK v3 databases prepared from published ISK application datasets are provided as Supplementary materials 1–3. Original second-generation ISK databases for testing conversion into ISK v3 are provided as Supplementary materials 4–6. A legacy FISK v2 database for testing import into AS-ISK v3 is provided as Supplementary material 7. TPS-ISK v2 databases for testing the Merge function are provided as Supplementary materials 8–12. Example outputs generated by ISK v3 are provided as Supplementary materials 13–18. A reviewer guide with source-publication information and suggested test workflows is provided as Supplementary material 19.

## Acknowledgements

We gratefully acknowledge Marina Piria, Zainab Al-Wazzan, Neil Angelo Abreo, Baran Yoğurtçuoğlu, Peter Barry, Hayrünisa Baş, Mahmood Bidarlord, Angela Boggero, Ratcha Chaichana, Xiaoyulong Chen, Udaya Priyantha Kankanamge Epa, Philippe Goulletquer, Gábor Herczeg, Jeffrey E. Hill, Wilson Iñiguez, Jenni Kakkonen, Qusaie Ebrahim Karam, Petra Kristan, Juliane Lukas, Levan Mumladze, Idrissa Ouédraogo, Dariusz Pietraszewski, Milica Ristovska, Predrag Simonović, Barbora Števove, Kristína Slovák Švolíková, Elena Tricarico, Hugo Verreycken, Hui Wei and Volodymyr Yuryshynets for post-release testing of ISK v3 and for providing valuable feedback. This study was conducted within the framework of a regionally coordinated programme on marine invasive species led by the Regional Organization for the Protection of the Marine Environment (ROPME). The views expressed in this article are those of the authors and do not necessarily reflect the official positions of ROPME or its Member States.

**Supplementary material 1**

**Sample AS-ISK v3 database derived from Vilizzi et al. (2026a)**

Authors: Lorenzo Vilizzi, Aisha Al-Marhoun

Data type: xlsx

Copyright notice: This dataset is made available under the Open Database License (http://opendatacommons.org/licenses/odbl/1.0/). The Open Database License (ODbL) is a license agreement intended to allow users to freely share, modify, and use this Dataset while maintaining this same freedom for others, provided that the original source and author are credited.

Link: to be assigned by the journal

**Supplementary material 2**

**Sample TAS-ISK v3 database derived from Vilizzi et al. (2022c)**

Authors: Lorenzo Vilizzi, Aisha Al-Marhoun

Data type: xlsx

Copyright notice: This dataset is made available under the Open Database License (http://opendatacommons.org/licenses/odbl/1.0/). The Open Database License (ODbL) is a license agreement intended to allow users to freely share, modify, and use this Dataset while maintaining this same freedom for others, provided that the original source and author are credited.

Link: to be assigned by the journal

**Supplementary material 3**

**Sample TPS-ISK v3 database derived from Vilizzi et al. (2024)**

Authors: Lorenzo Vilizzi, Aisha Al-Marhoun

Data type: xlsx

Copyright notice: This dataset is made available under the Open Database License (http://opendatacommons.org/licenses/odbl/1.0/). The Open Database License (ODbL) is a license agreement intended to allow users to freely share, modify, and use this Dataset while maintaining this same freedom for others, provided that the original source and author are credited.

Link: to be assigned by the journal

**Supplementary material 4**

**Original AS-ISK v2 database for conversion into ISK v3 from Vilizzi et al. (2026a)**

Authors: Lorenzo Vilizzi, Aisha Al-Marhoun

Data type: xlsx

Copyright notice: This dataset is made available under the Open Database License (http://opendatacommons.org/licenses/odbl/1.0/). The Open Database License (ODbL) is a license agreement intended to allow users to freely share, modify, and use this Dataset while maintaining this same freedom for others, provided that the original source and author are credited.

Link: to be assigned by the journal

**Supplementary material 5**

**Original TAS-ISK v2 database for conversion into ISK v3 from Vilizzi et al. (2022c)**

Authors: Lorenzo Vilizzi, Aisha Al-Marhoun

Data type: xlsx

Copyright notice: This dataset is made available under the Open Database License (http://opendatacommons.org/licenses/odbl/1.0/). The Open Database License (ODbL) is a license agreement intended to allow users to freely share, modify, and use this Dataset while maintaining this same freedom for others, provided that the original source and author are credited.

Link: to be assigned by the journal

**Supplementary material 6**

**Original TPS-ISK v2 database for conversion into ISK v3 from Vilizzi et al. (2024)**

Authors: Lorenzo Vilizzi, Aisha Al-Marhoun

Data type: xlsx

Copyright notice: This dataset is made available under the Open Database License (http://opendatacommons.org/licenses/odbl/1.0/). The Open Database License (ODbL) is a license agreement intended to allow users to freely share, modify, and use this Dataset while maintaining this same freedom for others, provided that the original source and author are credited.

Link: to be assigned by the journal

**Supplementary material 7**

**Legacy FISK v2 database for import into AS-ISK v3 from Vilizzi and Copp (2013)**

Authors: Lorenzo Vilizzi, Aisha Al-Marhoun

Data type: xls

Copyright notice: This dataset is made available under the Open Database License (http://opendatacommons.org/licenses/odbl/1.0/). The Open Database License (ODbL) is a license agreement intended to allow users to freely share, modify, and use this Dataset while maintaining this same freedom for others, provided that the original source and author are credited.

Link: to be assigned by the journal

**Supplementary material 8**

**TPS-ISK v2 merge-test database A from Vilizzi et al. (2024)**

Authors: Lorenzo Vilizzi, Aisha Al-Marhoun

Data type: xlsx

Copyright notice: This dataset is made available under the Open Database License (http://opendatacommons.org/licenses/odbl/1.0/). The Open Database License (ODbL) is a license agreement intended to allow users to freely share, modify, and use this Dataset while maintaining this same freedom for others, provided that the original source and author are credited.

Link: to be assigned by the journal

**Supplementary material 9**

**TPS-ISK v2 merge-test database B from Vilizzi et al. (2024)**

Authors: Lorenzo Vilizzi, Aisha Al-Marhoun

Data type: xlsx

Copyright notice: This dataset is made available under the Open Database License (http://opendatacommons.org/licenses/odbl/1.0/). The Open Database License (ODbL) is a license agreement intended to allow users to freely share, modify, and use this Dataset while maintaining this same freedom for others, provided that the original source and author are credited.

Link: to be assigned by the journal

**Supplementary material 10**

**TPS-ISK v2 merge-test database C from Vilizzi et al. (2024)**

Authors: Lorenzo Vilizzi, Aisha Al-Marhoun

Data type: xlsx

Copyright notice: This dataset is made available under the Open Database License (http://opendatacommons.org/licenses/odbl/1.0/). The Open Database License (ODbL) is a license agreement intended to allow users to freely share, modify, and use this Dataset while maintaining this same freedom for others, provided that the original source and author are credited.

Link: to be assigned by the journal

**Supplementary material 11**

>**TPS-ISK v2 merge-test database D from Vilizzi et al. (2024)**

Authors: Lorenzo Vilizzi, Aisha Al-Marhoun

Data type: xlsx

Copyright notice: This dataset is made available under the Open Database License (http://opendatacommons.org/licenses/odbl/1.0/). The Open Database License (ODbL) is a license agreement intended to allow users to freely share, modify, and use this Dataset while maintaining this same freedom for others, provided that the original source and author are credited.

Link: to be assigned by the journal

**Supplementary material 12**

**TPS-ISK v2 merge-test database E from Vilizzi et al. (2024)**

Authors: Lorenzo Vilizzi, Aisha Al-Marhoun

Data type: xlsx

Copyright notice: This dataset is made available under the Open Database License (http://opendatacommons.org/licenses/odbl/1.0/). The Open Database License (ODbL) is a license agreement intended to allow users to freely share, modify, and use this Dataset while maintaining this same freedom for others, provided that the original source and author are credited.

Link: to be assigned by the journal

**Supplementary material 13**

**ROC threshold report generated by ISK v3 from Vilizzi et al. (2021)**

Authors: Lorenzo Vilizzi, Aisha Al-Marhoun

Data type: pdf

Copyright notice: This dataset is made available under the Open Database License (http://opendatacommons.org/licenses/odbl/1.0/). The Open Database License (ODbL) is a license agreement intended to allow users to freely share, modify, and use this Dataset while maintaining this same freedom for others, provided that the original source and author are credited.

Link: to be assigned by the journal

**Supplementary material 14**

**Risk summary by species list generated by ISK v3 from Vilizzi et al. (2026a)**

Authors: Lorenzo Vilizzi, Aisha Al-Marhoun

Data type: xlsx

Copyright notice: This dataset is made available under the Open Database License (http://opendatacommons.org/licenses/odbl/1.0/). The Open Database License (ODbL) is a license agreement intended to allow users to freely share, modify, and use this Dataset while maintaining this same freedom for others, provided that the original source and author are credited.

Link: to be assigned by the journal

**Supplementary material 15**

**Risk summary by organismal group category generated by ISK v3 from Vilizzi et al. (2026a)**

Authors: Lorenzo Vilizzi, Aisha Al-Marhoun

Data type: xlsx

Copyright notice: This dataset is made available under the Open Database License (http://opendatacommons.org/licenses/odbl/1.0/). The Open Database License (ODbL) is a license agreement intended to allow users to freely share, modify, and use this Dataset while maintaining this same freedom for others, provided that the original source and author are credited.

Link: to be assigned by the journal

**Supplementary material 16**

**Single screening report generated by ISK v3 from Vilizzi et al. (2026a)**

Authors: Lorenzo Vilizzi, Aisha Al-Marhoun

Data type: xlsx

Copyright notice: This dataset is made available under the Open Database License (http://opendatacommons.org/licenses/odbl/1.0/). The Open Database License (ODbL) is a license agreement intended to allow users to freely share, modify, and use this Dataset while maintaining this same freedom for others, provided that the original source and author are credited.

Link: to be assigned by the journal

**Supplementary material 17**

**Grouped screening report generated by ISK v3 from Vilizzi et al. (2026a)**

Authors: Lorenzo Vilizzi, Aisha Al-Marhoun

Data type: xlsx

Copyright notice: This dataset is made available under the Open Database License (http://opendatacommons.org/licenses/odbl/1.0/). The Open Database License (ODbL) is a license agreement intended to allow users to freely share, modify, and use this Dataset while maintaining this same freedom for others, provided that the original source and author are credited.

Link: to be assigned by the journal

**Supplementary material 18**

**Compact exported database generated by ISK v3 from Vilizzi et al. (2026a)**

Authors: Lorenzo Vilizzi, Aisha Al-Marhoun

Data type: xlsx

Copyright notice: This dataset is made available under the Open Database License (http://opendatacommons.org/licenses/odbl/1.0/). The Open Database License (ODbL) is a license agreement intended to allow users to freely share, modify, and use this Dataset while maintaining this same freedom for others, provided that the original source and author are credited.

Link: to be assigned by the journal

**Supplementary material 19**

**Reviewer guide for sample databases and example outputs**

Authors: Lorenzo Vilizzi, Aisha Al-Marhoun

Data type: docx

Copyright notice: This document is made available under the Creative Commons Attribution License (CC BY 4.0), permitting use, distribution and reproduction provided that the original author and source are credited.

Link: to be assigned by the journal

## Notes

https://zenodo.org/doi/10.5281/zenodo.20414051

### Competing Interest Statement

The authors have declared no competing interest.

